# *De novo* generation of T-cell receptors with desired epitope-binding property by leveraging a pre-trained large language model

**DOI:** 10.1101/2023.10.18.562845

**Authors:** Jiannan Yang, Bing He, Yu Zhao, Feng Jiang, Zhonghuang Wang, Yixin Guo, Zhimeng Xu, Bo Yuan, Jiangning Song, Qingpeng Zhang, Jianhua Yao

## Abstract

Generating T-cell receptors (TCRs) with desired epitope-binding properties is a fundamental step in the development of immunotherapies, yet heavily relies on laborious and expensive wet experiments. Recent advancements in generative artificial intelligence have demonstrated promising power in protein design and engineering. In this regard, we propose a large language model, termed Epitope-Receptor-Transformer (ERTransformer), for the *de novo* generation of TCRs with the desired epitope-binding property. ERTransformer is built on EpitopeBERT and ReceptorBERT, which are trained using 1.9 million epitope sequences and 33.1 million TCR sequences, respectively. To demonstrate the model capability, we generate 1000 TCRs for each of the five epitopes with known natural TCRs. The artificial TCRs exhibit low sequence identity (average Bit-score 27.64 with a standard deviation of 1.50) but high biological function similarity (average BLOSUM62 score 32.32 with a standard deviation of 12.01) to natural TCRs. Furthermore, the artificial TCRs are not very structurally identical to natural ones (average RMSD 2.84 Å with a standard deviation of 1.21 Å) but exhibit a comparable binding affinity towards the corresponding epitopes. Our work highlights the tremendous potential of applying ERTransformer to generate novel TCRs with desired epitope-binding ability.

## 1. Introduction

Large language models experienced extraordinary advances in content generation in recent years. The ChatGPT, which was built on top of the Generative Pre-trained Transformer 3 (GPT-3) family of language models, has shown promising power in dialogue generation. Protein sequences are analogous to human languages. These amino acids (“letters”) arrange to form secondary structural elements (“words”), which assemble to form domains (“sentences”) that perform a biological function (“meaning”). There are some preliminary studies to generate artificial proteins using large language models. ProtGPT2, a language model trained on the protein space, is capable of generating *de novo* proteins with topologies not present in existing structure databases ^1^. ProGen generated artificial proteins with similar catalytic efficiencies to natural lysozymes ^2^. ProteinMPNN generates protein sequences from its structure ^3^ and facilitates the de novo design of luciferases ^4^. However, these models are still limited to generating functionally similar artificial proteins from the known proteins themselves. In living organisms, proteins interact with each other to collectively perform biological functions ^5^. Compared to generating functionally similar proteins, generating functional interacting partner proteins for a given protein is a more challenging task for language models, but is also more essential for biomedicine and bioengineering applications. To demonstrate the feasibility of such generation, we took the interacting proteins in a basic biological process, the cell-mediated immune response, as an example to perform a proof of concept study.

Upon exposure to an antigen, which may originate from a pathogen or a vaccination, the cell-mediated immune response is activated by T cells ^6^. This process involves the recognition of an epitope, a specific part of the antigen presented by a major histocompatibility complex (MHC), by the T cell receptor (TCR). The binding between an epitope and T cell receptors plays a critical role in the activity and specificity of T cells ^7^. For example, adoptive cell therapy (ACT) relies on TCRs to redirect T cells, because its TCRs can recognize epitopes present by MHC on the cell membrane of the specific cancer cells. ACT showed promising potential for cancer therapy in recent trials ^8–10^. Recently, the first TCR therapy, termed KIMMTRAK, was approved by FDA for the treatment of metastatic uveal melanoma ^11^. Understanding the characteristics of epitopes, TCRs, and their interactions is critical to developing effective immune therapies.

Despite the efficacy of TCR-based immune therapy, developing such therapy for a patient is currently laborious and too expensive to afford ^12^. One challenge is to identify tumor-reactive TCRs. The commonly-used, standardized approach is to isolate a tumor-reactive T cell, which relies on a proper source that ideally harbors a high avidity TCR and reaches a significant level of frequency and purity ^13^. This approach is challenging, especially for poorly immunogenic tumors ^14^. After retrieving tumor-reactive T cells, successfully identifying tumor-reactive TCR sequences remains a great challenge, which requires profiling the diversity of millions of TCR molecules in the retrieved samples. Thus, *in silico* approaches that could directly generate the tumor-reactive TCRs given specific epitopes of tumor cells would greatly speed up the immune therapy design.

With the advances in high-throughput sequencing technologies, the amount of known epitopes and TCR sequences is increasing dramatically, leading to the emergence of various computational models for identifying tumor-reactive TCRs. Classic models such as GLIPH ^15,16^, and TCRMatch ^17^ utilize the sequence motifs of TCRs to predict the binding affinity between TCRs and antigens. Recently, the birth of Bidirectional Encoder Representations from Transformers (BERT) models revolutionized the natural language processing (NLP) field ^18^. Because of the similarities between protein sequences and language sentences, several studies applied BERT models to TCR-related tasks ^19,20^. TCR-BERT ^19^ utilized over 8,000 TCR sequences on the self-supervised learning tasks, and showed promising performance on the downstream tasks, such as TCR-antigen binding affinity prediction and sequence clustering. Han et al. ^20^ presented a BERT-based model by fine-tuning the pre-trained Tasks Assessing Protein Embeddings (TAPE) ^21^ model to predict SARS-CoV-2 T-cell epitope-specific TCR recognition. However, to the best of our knowledge, all the existing approaches only focused on the modeling of natural TCR sequences. While the adaptive immune system relies on the cooperative interaction of both epitopes and TCRs, the absence of epitope modeling will significantly hinder our understanding of the immune system using computational models. In addition, compared with tens of billions of language data utilized in NLP areas, the data on TCR-related tasks, especially the data about epitope-TCR binding pairs are still relatively small. Thus, none of the existing approaches have attempted to tackle the challenge of sequence generation in this domain, which limits the potential of computational models for generating TCR sequences with desired properties.

To address the aforementioned limitations, we propose a large language model Epitope-Receptor-Transformer (ERTransformer) for the artificial *de novo* generation of TCR with desired epitope-binding properties. To develop ERTransformer, we first built ER-BERT, which is composed of two BERT modules: the EpitopeBERT and the ReceptorBERT. Both modules are pre-trained on a large amount of epitope (EpitopeBERT) and receptor (ReceptorBERT) sequences to learn the common rules in these sequences. Subsequently, we created the ERTransformer model in which the encoder and decoder blocks are derived from the pre-trained EpitopeBERT and ReceptorBERT, respectively. To capture the binding rules, we combined the ER-BERT with a multilayer perceptron (MLP) head layer and fine-tuned it on the Binding Specificity Prediction (BSP) task (ER-BERT-BSP), which can be used to determine the quality of the generated TCR sequences. We introduced ERTransformer to generate the complementarity-determining region 3 (CDR-3) part of TCR beta (TRB) chain sequences and verified the quality of the generated TRB sequences using three tools, including external discriminators for the determination of binding specificity, the Basic Local Alignment Search Tool (BLAST) ^22^ for the sequence identity, and BLOcks SUbstition Matrix (BLOSUM62) matrix^23^ for the biological function similarity. The results demonstrate that ERTransformer can generate TRBs with good binding specificity to the corresponding epitopes. The artificially generated TRBs are not similar in sequence to the natural TRBs, as demonstrated by an average Bit-score of 27.64 with a standard deviation (std) of 1.50 with BLAST. Generally, a Bit-score of lower than 40 indicates low sequence similarity ^24^. However, the artificial TRBs have a high biological function similarity with natural TRBs, which is demonstrated by an average BLOSUM62 score of 32.32 with std 12.01. A positive BLOSUM62 score indicates the presence of biological function similarity, which increases with the increasing value of the score ^23^. We further utilized AlphaFold2 ^25^, TCRDock ^26^ and RosettaDock ^27^ to estimate the structure and binding affinity of the artificially generated TCRs. Our analysis revealed that the artificial TRBs are not structurally identical to natural ones (average RMSD 2.84 Å with a standard deviation of 1.21 Å), but exhibit a comparable binding affinity towards corresponding epitopes. In addition, the results show that ERTransformer can capture the key amino acids that determine the docking positions of epitopes and TCRs using the integrated gradients method ^28^.

## 2. Results

### 2.1 The framework of ERTransformer

The training framework of the Epitope-Receptor-Transformer (ERTransformer) is illustrated in Figure 1(A-E). ERTransformer is built on the ER-BERT, which consists of pre-trained EpitopeBERT and ReceptorBERT, both of which use the standard Bidirectional Encoder Representations from Transformers (BERT) architecture ^18^. The BERT language model ^18^ was originally proposed for natural language processing tasks, such as translation ^29^ and sentiment analysis ^30^. Both EpitopeBERT and ReceptorBERT have 12 encoder blocks, each of which consists of two layers: the multi-head self-attention layer and the feed-forward layer. The architecture of stack encoder blocks allows the BERT model to learn the high-dimensional and complex interactions between tokens to capture the “grammar” rules in the input sequences. The token is the basic unit to form the input and output sequences. We hypothesize that Epitope and TCR sequences can be represented as a series of amino acid sequences composed of single-letter symbols, such as epitope “ATDALMTGY”, which is very similar to language sentences. The similarities of amino acid sequences and language sentences allow us to naturally introduce the BERT model to capture the rules in the epitope and TCR sequences.

**Figure 1.**
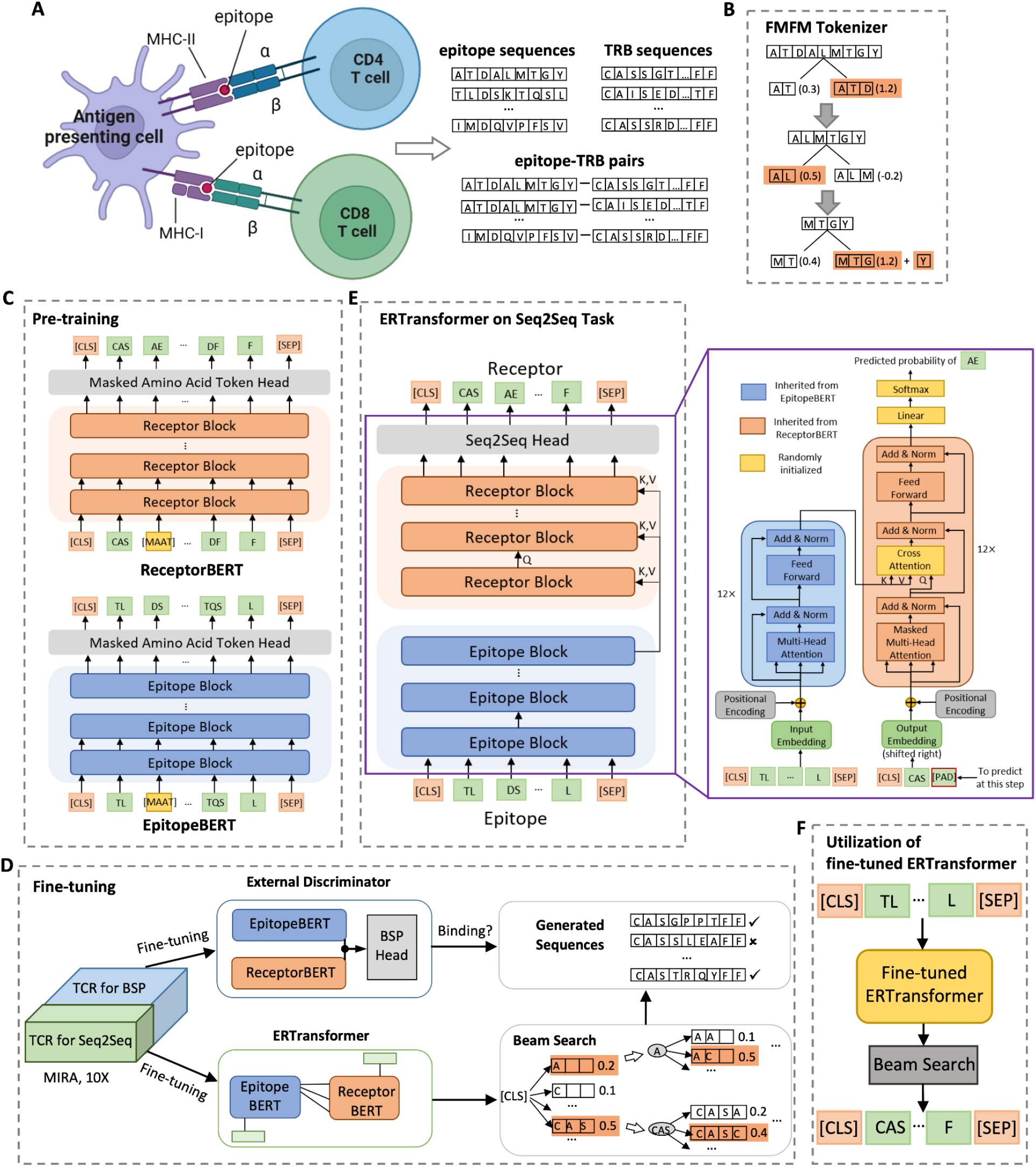
The framework of Epitope-Receptor-Transformer (ERTransformer). **(A)** We collected 1,929,016 epitope sequences, 33,088,640 TRB sequences, and 176,268 epitope-TRB pairs, considering both CD8 and CD4 T cells. **(B)** The proposed FMFM tokenizer separates an amino acid sequence by maximizing the frequencies of the motif. The example shows that the epitope sequence ATDALMTGY is divided into 4 tokens: “ATD”, “AL”, “MTG”, and “Y”. **(C)** The pre-training of ER-BERT consists of two parts: EpitopeBERT and ReceptorBERT are pre-trained on the Masked Amino Acid Token (MAAT) task in a self-supervised manner using collected epitope sequences and TRB sequences, respectively. **(D)** The fine-tuning of the ERTransformer in this study to generate epitope-binding TCRs. Given an external dataset such as MIRA or 10X that contains epitope-TCR pairs, for each epitope, we used 20% of its binding TCRs to fine-tune the ERTransformer, which consists of the pre-trained EpitopeBERT and ReceptorBERT. Then we utilized the Beam Search method to generate TCR sequences. The remaining 80% of the binding TCRs were used to fine-tune the ER-BERT-BSP, which consists of the pre-trained EpitopeBERT and ReceptorBERT and a Binding Specificity Prediction (BSP) head. The ER-BERT-BSP acts as an external discriminator to determine whether the generated TCRs can bind to the given epitope. **(E)** The architecture of ERTransformer for the sequence generation (Seq2seq task) from the epitope sequence to the receptor sequence. ERTransformer consists of 12 epitope blocks (encoder blocks) and 12 receptor blocks (decoder blocks) derived from pre-trained EpitopeBERT and ReceptorBERT, respectively. In each epitope block, the parameters of all layers are inherited from EpitopeBERT. In each receptor block, the parameters of the cross-attention layer are initialized randomly, and the parameters of all other layers are inherited from ReceptorBERT. The *key* (*Q_E_*) matrix, *value* (*K_E_*) matrix, and *query* (*V_R_*) matrix are the input of the cross-attention layer. The *Q_E_* and *K_E_* are the embeddings of the epitope, while *V*_*R*_ are the embeddings of the amino acid sequences that were already generated for the TCR. **(F)** The application of the fine-tuned ERTransformer to generate TCR for a new epitope.

To train EpitopeBERT and ReceptorBERT, we collected a comprehensive dataset containing 1,929,016 epitope sequences, 33,088,640 TRB sequences, and 176,268 epitope-TRB pairs from ten public and two in-house datasets (see Methods and Datasets for more details), considering both CD8 and CD4 T cells (Figure 1(A)). Since the amount of TRA sequences (408,722) was far smaller than that of TRB sequences (33,088,640), we focused on TRB in this study. Given the input epitope or TRB sequence with a length *l*, after tokenization using a tokenizer, the input is separated into *M* tokens. Different from English sentences with words separated by spaces naturally, epitope and TCR sequences are amino acid sequences without natural separations. We proposed a new tokenizer named the Forward Maximum Frequency Matching (FMFM) tokenizer (Figure 1(B)), which could separate an amino acid sequence into tokens by maximizing the frequencies of the motif. The motif herein is defined as the pattern of amino acid combinations that are recurrently observed in a protein. Motifs are widely present in many proteins and peptides ^31^, and their frequencies are highly associated with distinct functions ^32^. In addition, for comparison, we introduced a commonly-used tokenizer that treats each amino acid as a token, which we named the unique amino acid (UAA) tokenizer. The generated tokens are then fed into the embedding layer. We then encoded the sequence embeddings with the BERT model to a “head” layer for pre-training or downstream tasks.

The TCR generation in this study has two major steps: pre-training and fine-tuning. First, we constructed ER-BERT with self-supervised pre-training of EpitopeBERT and ReceptorBERT on the Masked Amino Acid Token (MAAT) task to learn the basic patterns that existed in epitope and TCR sequences using the above-mentioned datasets (Figure 1(C)). Second, we constructed ERTransformer with parameters from ER-BERT and fine-tuned it to generate TCR sequences (Figure 1(D)). We used two external datasets (see Methods) to validate ERTransformer’s efficacy in generating TCR sequences. Specifically, given an external dataset, we used 20% of binding TCR sequences for each epitope to fine-tune the ERTransformer on the Seq2Seq task (from epitope sequence to generate TCR sequence). ERTransformer consists of 12 epitope blocks (encoder blocks) and 12 receptor blocks (decoder blocks) derived from pre-trained EpitopeBERT and ReceptorBERT, respectively. As illustrated in Figure 1(E), in each receptor block, only the parameters of the cross-attention layer are initialized randomly, the parameters of all other layers are inherited from ReceptorBERT. Likewise, the parameters of all layers in the epitope block are inherited from EpitopeBERT. During the training of the Seq2Seq task, the epitope block will be fed the whole epitope sequence to produce the epitope embedding, which is the high-dimensional latent representation of the epitope. The receptor block is responsible for generating the target token (amino acids) of TCR step by step. At each step, the receptor block receives epitope embedding through the cross-attention layer. Besides the epitope embedding, the other input to the receptor block is the amino acids that were already generated for the TCR. Then, after the fine-tuning of ERTransformer, we utilized Beam Search ^33^, a commonly-used language generation method, to generate the TCR sequences for each epitope. The remaining 80% TCR sequences were used to train the ER-BERT-BSP, which is the pre-trained ER-BERT with a multilayer perceptron (MLP) head, on the Binding Specificity Prediction (BSP) task as an external discriminator to determine whether the generated TCRs could bind to the given epitopes. The ER-BERT-BSP uses ER-BERT to learn the “binding rule” of epitopes and TCRs (Figure 1(D)). Specifically, given one epitope-TCR pair, we utilized the pre-trained EpitopeBERT and ReceptorBERT to obtain the learned representations and appended a “BSP” head to determine whether the TCR binds to the given epitope. To generate binding TCR sequences for a new epitope, as shown in Figure 1(F), we could use the fine-tuned ERTransformer through the Beam Search method directly. For more details, please refer to the Methods and Datasets.

### 2.2 ER-BERT achieved superior performance in clustering epitope-specific TCRs

ER-BERT aims to learn the high-dimensional latent representations of epitopes and TCRs to further capture their complex relationships. To validate whether the learned representations are biologically meaningful, we used the Principal Component Analysis (PCA) and t-distributed stochastic neighbor embedding (t-SNE)^34^ to reduce the latent representations’ dimensions and obtain the visualizations of these representations. We then further investigated whether the representations of epitope-specific TCRs could cluster together and computed the Davies-Bouldin index (DBI) as a quantitative metric to measure the effects of clustering (the smaller the DBI is, the better the clustering performance). We also used TCRBert to compare the clustering effects with ER-BERT.

We reported the DBI scores of all the epitopes and their corresponding TRBs with random seed settings in Figure 2(D). We found that the ER-BERT with the FMFM tokenizer trained after the Seq2Seq task (fine-tuned ERTransformer) achieved the best DBI score (mean 10.27, std 5.66), while this model trained after the MAAT task (pre-trained ER-BERT) also achieved the worst DBI score (mean = 26.88, std = 9.65). Among the three tasks, i.e. MAAT task (pre-trained ER-BERT), BSP task (fine-tuned ER-BERT-BSP), and Seq2seq task (fine-tuned ERTransformer), we found that ER-BERT with two tokenizers both achieved the best clustering effects after being trained on the Seq2Seq task (fine-tuned ERTransformer). It’s noted that for the Seq2Seq task (fine-tuned ERTransformer), the ER-BERT model with FMFM tokenizer has better DBI performance than the one with UAA tokenizer. In general, TCRBert’s performance (mean = 20.86, std = 7.56) was worse than that of ER-BERT with both two tokenizers. In conclusion, ER-BERT can provide superior epitope-specific clustering compared with the state-of-the-art TCRBert. ER-BERT performs the best using the FMFM tokenizer trained after the Seq2Seq task (fine-tuned ERTransformer).

**Figure 2.**
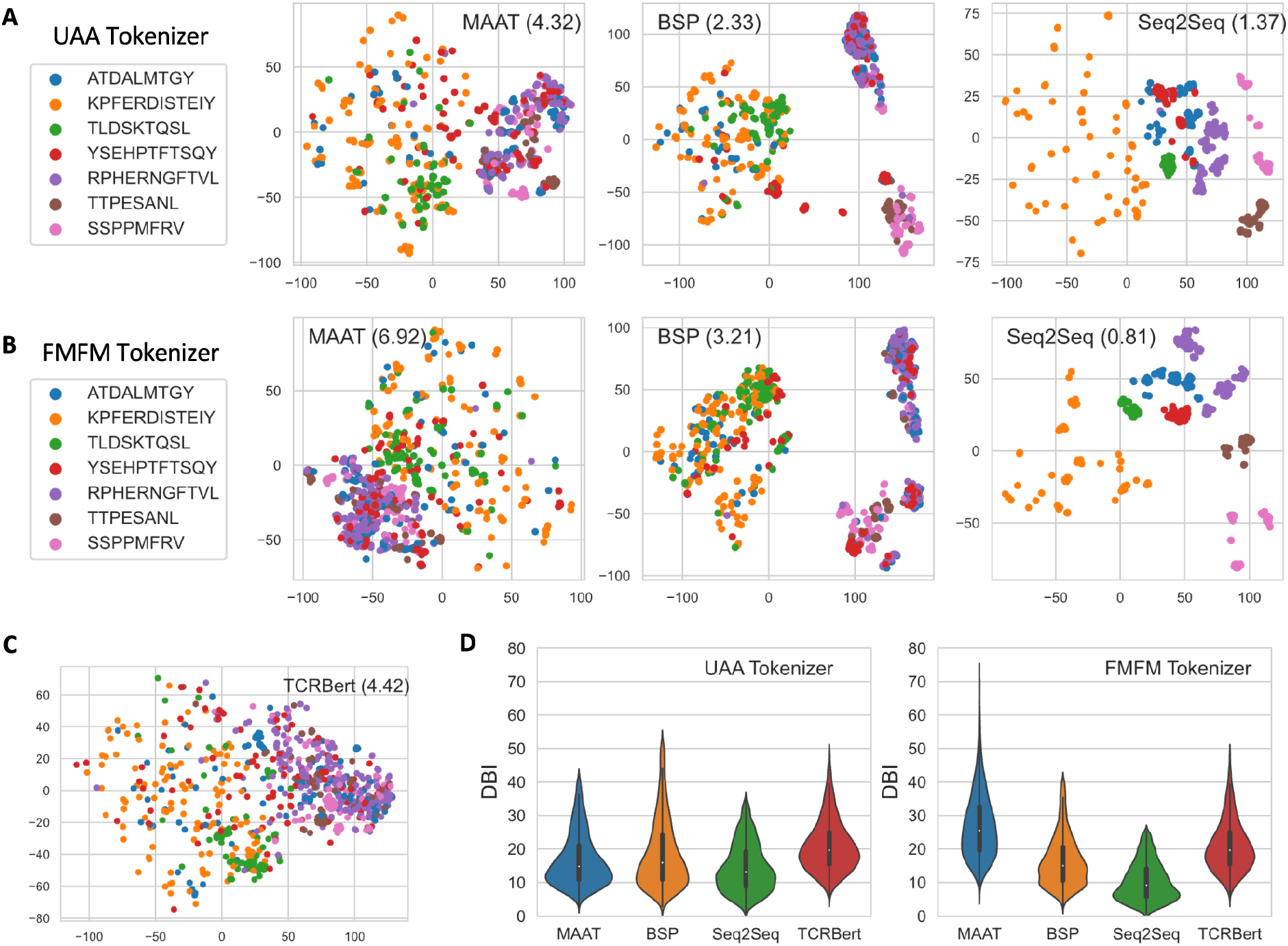
The clustering of epitope-specific TCRs using the representations generated by ER-BERT with two tokenizers and TCRBert. We selected seven epitopes from different viruses as an example to present the clustering ability of ER-BERT using the PCA and t-SNE methods. The values in each sub-figure denote the DBI scores. (A), (B) and (C) show the clustering of epitope-specific TCRs using ER-BERT with the UAA tokenizer (A), ER-BERT with the FMFM tokenizer (B), and TCRBert (C), respectively. (D) shows the DBI scores considering all the epitopes and their corresponding TRBs with random seed settings of ER-BERT with two tokenizers after training on three tasks, and TCRBert.

We selected seven epitopes that belong to Hepatitis C virus (“ATDALMTGY”), SARS-CoV-2 (“KPFERDISTEIY”, “TLDSKTQSL”), Human herpesvirus 5 (“YSEHPTFTSQY”, “RPHERNGFTVL”), TL8 of transcription activator (“TTPESANL”), and Murid herpesvirus 1 (“SSPPMFRV”) and their corresponding TRBs as an example. These epitopes cover those presented by MHC Class-I (“ATDALMTGY”: HLA-A*01:01, “YSEHPTFTSQY”: HLA-A*01:01, “TTPESANL”: Mamu-A1*001:01, “SSPPMFRV”: H2-Kb) and Class-II (“KPFERDISTEIY”: HLA-B∗40:01, “TLDSKTQSL”: HLA-B*08:01, “RPHERNGFTVL”: HLA-B*07:02). The 2-dimensional visualization of the representations of paired epitopes and TCRs using the t-SNE method is shown in Figure 2(A-C). We found that the clustering effects of ER-BERT with two tokenizers and TCRBert are consistent with the general performance on all the epitopes. And the clustering performance of TCRBert is still in general worse than ER-BERT with two tokenizers trained after the three tasks.

### 2.3 ERTransformer can generate artificial TCRs with desired epitope-binding specificity

Given one epitope, fine-tuned ERTransformer with parameters from ER-BERT could generate thousands of TCRs. The important issue is how to determine the quality of generated sequences. Here, we trained two external discriminators, including ER-BERT-BSP and DeepTCR ^35^, to evaluate the ability of the generated TCRs to bind to input epitopes. Given one dataset, such as MIRA and 10X, we used 80% data to train the external discriminators and the remaining 20% of the data to fine-tune the ERTransformer for the TCR generation task. ER-BERT-BSP is an extension of the ER-BERT model, incorporating a multilayer perceptron (MLP) head that has been specifically trained on the BSP task. In this study, the model DeepTCR ^35^, originally developed by Sidhom et al., was retrained utilizing the identical dataset as ER-BERT-BSP due to the unavailability of a pre-trained version (see Methods for more details). The performance of these two discriminators on MIRA and 10X datasets are reported in Figure 3(A, B). ER-BERT-BSP achieved a ROC-AUC of 0.90 and 0.86 on MIRA and 10X, respectively, which significantly outperforms DeepTCR (ROC-AUC of 0.77 and 0.76, respectively), indicating the superior performance of ER-BERT-BSP on the binding specificity prediction task.

**Figure 3.**
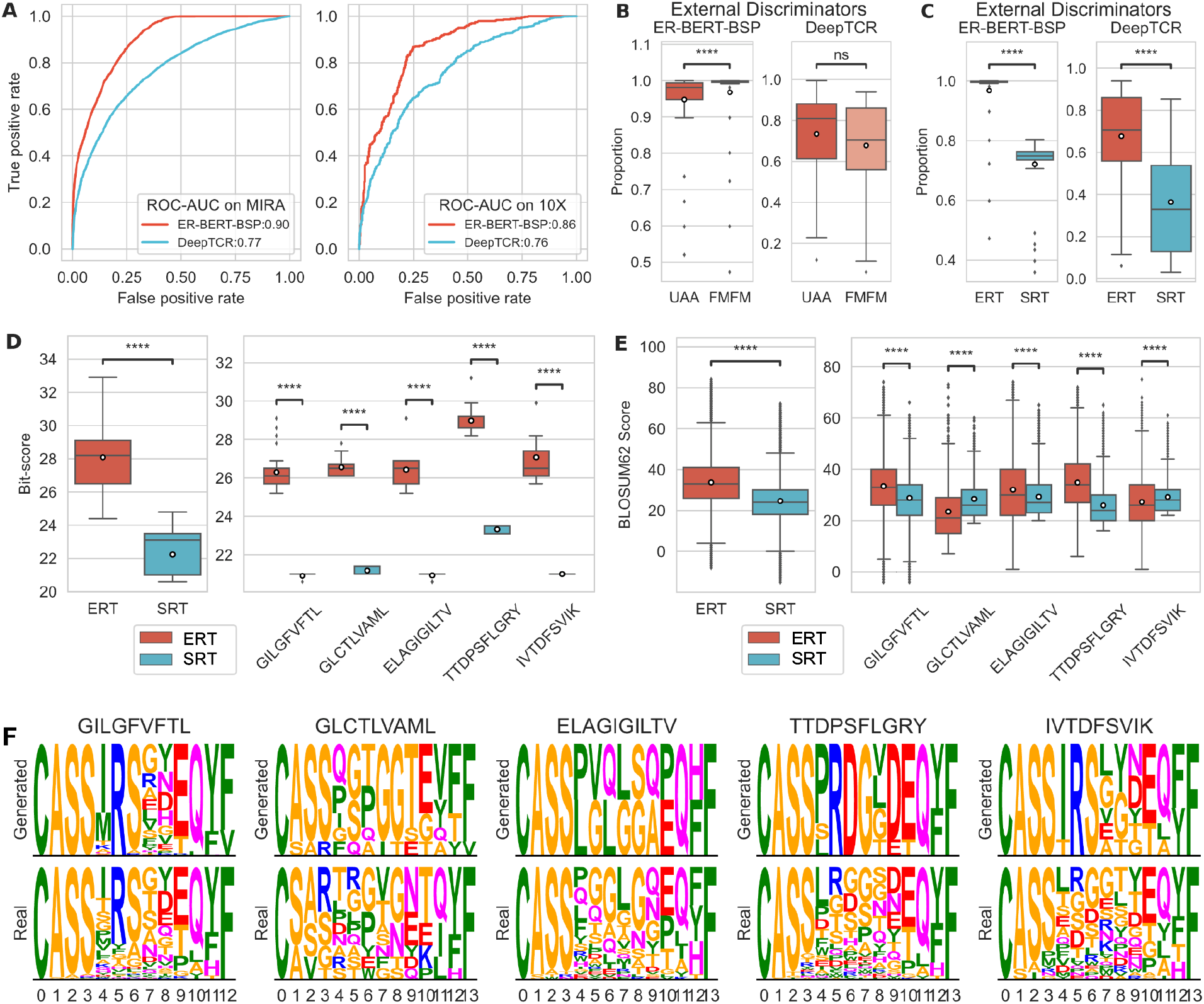
The performance of ER-BERT on the TCR generation task. (A) The ROC-AUC curves of external discriminators on the MIRA (left) and 10X (right) datasets. For one dataset, we used 80% of the data to train the external discriminators and the remaining 20% of the data to fine-tune the ERTransformer (ERT) and Semi-Random-Transformer (SRT) for the TCR generation task. (B-C) The proportion of the generated TRBs that are determined as binding to the given epitopes by the external discriminator ER-BERT-BSP (left) and DeepTCR (right). In (B), TRBs are generated by ERTransformer using the UAA and FMFM tokenizer, respectively. FMFM tokenizer showed superior performance compared to UAA tokenizer in the TRB generation task. Thus, FMFM was used as the default tokenizer for ERTransformer in the following analysis. In (C), TRBs are generated by ERTransformer (ERT) and Semi-Random-Transformer (SRT), respectively. (D) The Bit-scores of BLAST by comparing the generated and natural TRBs. (E) The BLOSUM62 scores by comparing the generated and natural TRBs. Note that in both (D) and (E), the left plot presents the metrics of all the epitopes that existed in the MIRA and 10X datasets, and the right plot shows the metrics for selected epitopes. The p-value annotation legend in (B-E) are: *ns*: *p* ≤ 1; *: 0. 01 < *p* ≤ 0. 05; **: 0. 001 < *p* ≤ 0. 01; ***: 0. 0001 < *p* ≤ 0. 001; ****: *p* ≤ 0. 0001. ERT is the abbreviation of ERTransformer. SRT is the abbreviation of Semi-Random-Transformer. The horizontal lines and circles in each box in (B-E) represent the median and mean values. (F) The sequence logo plots of the selected five epitopes, where the upper and lower sub-plots display the sequence logo plot of the generated and corresponding natural TRBs with the same length, respectively.

To identify the optimal tokenizer method for ERTransformer, we utilized the fine-tuned ERTransformer with two tokenizers to generate 1,000 TRB sequences for each epitope in the MIRA and 10X datasets. Then we used the two external discriminators to evaluate the quality of the generated TRBs. Note that we have removed the generated TRB sequences that also exist in the dataset for the fine-tuning in 10X or MIRA (Table S1 in Supplementary Information). Figure 3(B) presents the proportions of the number of binding TRBs determined by the two external discriminators separately. When using ER-BERT-BSP as the external discriminator (Figure 3(B) left), more than 90% of the TRBs generated by ERTransformer with two tokenizers were determined as binding to the input epitopes. Among the two tokenizers, the FMFM tokenizer performed significantly better (*p* ≤ 0. 0001) than the UAA tokenizer in the TRB generation task (Figure 3(B) left). When using DeepTCR as the external discriminator, the performance of ERTransformer with the UAA tokenizer and FMFM tokenizer showed no significant difference (Figure 3(B) right). Thus, FMFM was used as the default tokenizer for ERTransformer in the following analysis (ERTransformer with FMFM tokenizer is abbreviated as ERT in Figure 3).

Because there is no existing computational approach to the TCR sequence generation task, we need to create a reasonable baseline to validate the performance of ER-BERT. Therefore, we created a baseline termed Semi-Random-Transformer (abbreviated as SRT in Figure 3) using the same architecture as ERTransformer but a randomly initialized BERT instead of EpitopeBERT and a previously published TCRBert ^19^ instead of the ReceptorBERT. We applied the method of generating TCR sequences using ERTransformer and the Semi-Random-Transformer fine-tuned on the same dataset (Method 4.3 and 4.5.3). For each epitope in the MIRA and 10X datasets, we applied ERTransformer and Semi-Random-Transformer fine-tuned on each dataset to generate 1,000 TRB sequences for the given epitope and then used the two external discriminators to evaluate the quality of the generated TRBs. Figure 3(C) presents the proportions of the number of binding TRBs determined by the two external discriminators separately. When using ER-BERT-BSP as the external discriminator (Figure 3(C) left), less than 75% of the TRBs generated by the Semi-Random-Transformer were determined as binding to the given epitopes, which was significantly lower than that of ERTransformer (*p* ≤ 0. 0001). When using DeepTCR as the external discriminator (Figure 3(C) right), the performance of the Semi-Random-Transformer dropped dramatically, with only 36% of the generated TRBs determined as binding to the given epitopes. Meanwhile, the performance of ERTransformer also dropped, but was still significantly better than the performance of the Semi-Random-Transformer (*p* ≤ 0. 0001). Combining these results, we conclude that the proposed ERTransformer framework shows great potential in the TCR sequence generation task. Even though TCRBert is not specifically designed for the sequence generation task, the Semi-Random-Transformer still shows great generalization power in the TCR sequence generation task after fine-tuning. In addition, with the pre-training and fine-tuning procedure, as well as the specially-designed FMFM tokenizer, ER-BERT can generate TCRs with great binding specificity (more TCRs that could bind to the given epitope).

### 2.4 Artificial TCRs mimic natural functionality despite sequence differences

To check whether the TCRs generated by ERTransformer are similar to the natural TCRs by their sequences, we applied two methods - The Basic Local Alignment Search Tool (BLAST) ^22^ and BLOSUM62 matrix^23^ to compare all the generated and natural TCRs. In addition, we also selected five epitopes from MIRA and 10X datasets. These epitopes are presented by HLA Class-I and belong to the Influenza A virus (“GILGFVFTL”: HLA-A*0201), SARS-CoV-2 virus (“TTDPSFLGRY”: HLA-A*01:01), Human herpesvirus 4 (“GLCTLVAML”: HLA-A*02:01, “IVTDFSVIK”: HLA-A*11:01), and Melan-A (“ELAGIGILTV”: HLA-A*02:01). These five epitopes have the most binding TCRs in the MIRA and 10X datasets, and also contain the full-length information of the TCRs which can be used for the structure estimation for the following analysis.

Given one generated TCR sequence, the BLAST tool is used to compare its amino acid sequence and calculate the statistical significance of the matched reference sequence. Bit-score is used to determine the sequence similarity in the BLAST tool, and the higher the bit-score, the better the sequence similarity. As shown in Figure 3(D), ERTransformer achieved an average Bit-score of 27.64 with std 1.50. Compared to the performance of the Semi-Random-Transformer, the Bit-score of the Semi-Random-Transformer (mean 22.04 with std 1.19) was significantly smaller than ERTransformer (*p* ≤ 0. 0001). The Bit-scores of the selected five epitopes also followed the same patterns, with ERTransformer achieving the better bit-scores than Semi-Random-Transformer. It’s known that a Bit-score of lower than 40 indicates a low sequence similarity ^24^. Thus, the artificial TCRs generated by both methods are not similar to the natural ones in their sequences.

The BLOSUM62 matrix^23^ provides a quantitative approach that determines whether an amino acid substitution is biologically conservative or nonconservative. The biologically conservative substitution, which indicates the presence of biological function similarity, has positive scores in the BLOSUM62 matrix. The biological function similarity between two sequences increases with the value of the positive BLOSUM62 score. In contrast, the nonconservative substitution, which indicates no biological function similarity, has negative scores in the BLOSUM62 matrix. Here, for each epitope, we computed the average BLOSUM62 scores of all the amino acid positions for all its generated and natural TRBs. As shown in Figure 3(E), ERTransformer achieved an average BLOSUM62 score of 32.32 with std 12.01, which was significantly larger (*p* < 0. 0001) than Semi-Random-Transformer (mean = 28.25 and std = 8.44). It reveals that, although artificial TCRs generated by both methods have biological function similarity to the natural ones, the TCRs generated by ERTransformer are significantly better than the ones by the baseline method.

We took a careful look at the sequence conservation of amino acids of the generated and natural TCRs, as shown in the sequence logo plots (Figure 3(F)) of the selected five epitopes. The TRB sequences were generated by ERTransformer using the FMFM tokenizer since this tokenizer achieves the best performance using the three tools mentioned above. Specifically, for one epitope, given all its corresponding natural TCRs, we first acquired the most common length of these TRB sequences. Then, we only kept the generated and natural TRBs with that length to further get the sequence logo plot. From the sequence logo plots in Figure 3(F), we found that the artificially generated TRBs have already captured the patterns of the natural TRBs. For example, most TRBs start with motif “CASS” and end with motif “FF” or “YF”, which is consistent with previous studies ^36^. Meanwhile, we found that the natural TRBs are more diverse compared to the artificial ones, especially considering the middle positions. This is probably due to that the artificial TRB sequences are generated following the epitope-TCR binding rules learned by the model, instead, the natural ones are generated by the process of combining random generation and natural selection ^37^.

In conclusion, ERTransformer already has the ability to capture the general patterns existing in the natural TCR sequences and could generate TCR sequences that are similar in biological function but not similar in sequence to the natural ones.

### 2.5 The artificial TCRs exhibit a binding affinity towards epitopes that closely resembles that of natural ones

In this section, we further investigated the quality of the generated TCRs in terms of structural and functional similarity to natural ones. We selected three epitopes “ELAGIGILTV” (PDB ID: 3hg1 ^38^, HLA-A*02:01), “GILGFVFTL” (PDB ID: 1oga ^39^, HLA-A*0201), “GLCTLVAML” (PDB ID: 3o4l ^40^, HLA-A*02:01) and their corresponding natural TRBs CDR3 “CAWSETGLGTGELFF” (PDB ID: 3hg1), “CASSSRSSYEOYF” (PDB ID: 1oga), “CSARDGTGNGYTF” (PDB ID: 3o4l) for which experimentally validated structures of the pMHC-TCR binding complex are available at Protein Data Bank (PDB). Given an artificial TRB (CDR3 part) generated by ERTransformer with the FMFM tokenizer for each of the above epitope, we retrieved the sequence of the corresponding full length natural TRB from the PDB structure and replaced the original CDR3 part with the artificial one to construct a full length artificial TRB. Then, we introduced TCRdock ^26^, which is a Alphafold-Multimer model that fine-tuned for TCR structure prediciton, to estimate the structure of the artificial TCR and used PyMOL to visualize the structure. The structure similarity between natural TCR and the artificial one is measured by the root-mean-square deviation (RMSD) of atomic positions. Figure 4(A) depicts the RMSD of artificial TRBs, specifically focusing on the CDR3 region, in comparison to the structure of natural ones. These artificial TRBs were selected from the top 100 generated TRBs based on their binding probabilities, as predicted by the ER-BERT-BSP model. Additionally, any generated TRBs with identical sequences to natural ones were removed from the selection. As presented in Figure 4(A), the generated artificial TRBs exhibit an average RMSD of 2.84 Å with a standard deviation of 1.21 Å for all three epitopes (3hg1: 2.23 ±1.02 Å; 1oga: 3.26 ± 1.31 Å; 3o4l: 3.22 ± 0.87 Å). In addition, the top three artificial TRBs predicted by the ER-BERT-BSP did not exhibit the smallest RMSD values, suggesting that these artificial TRBs are not the most structurally identical to natural ones.

**Figure 4.**
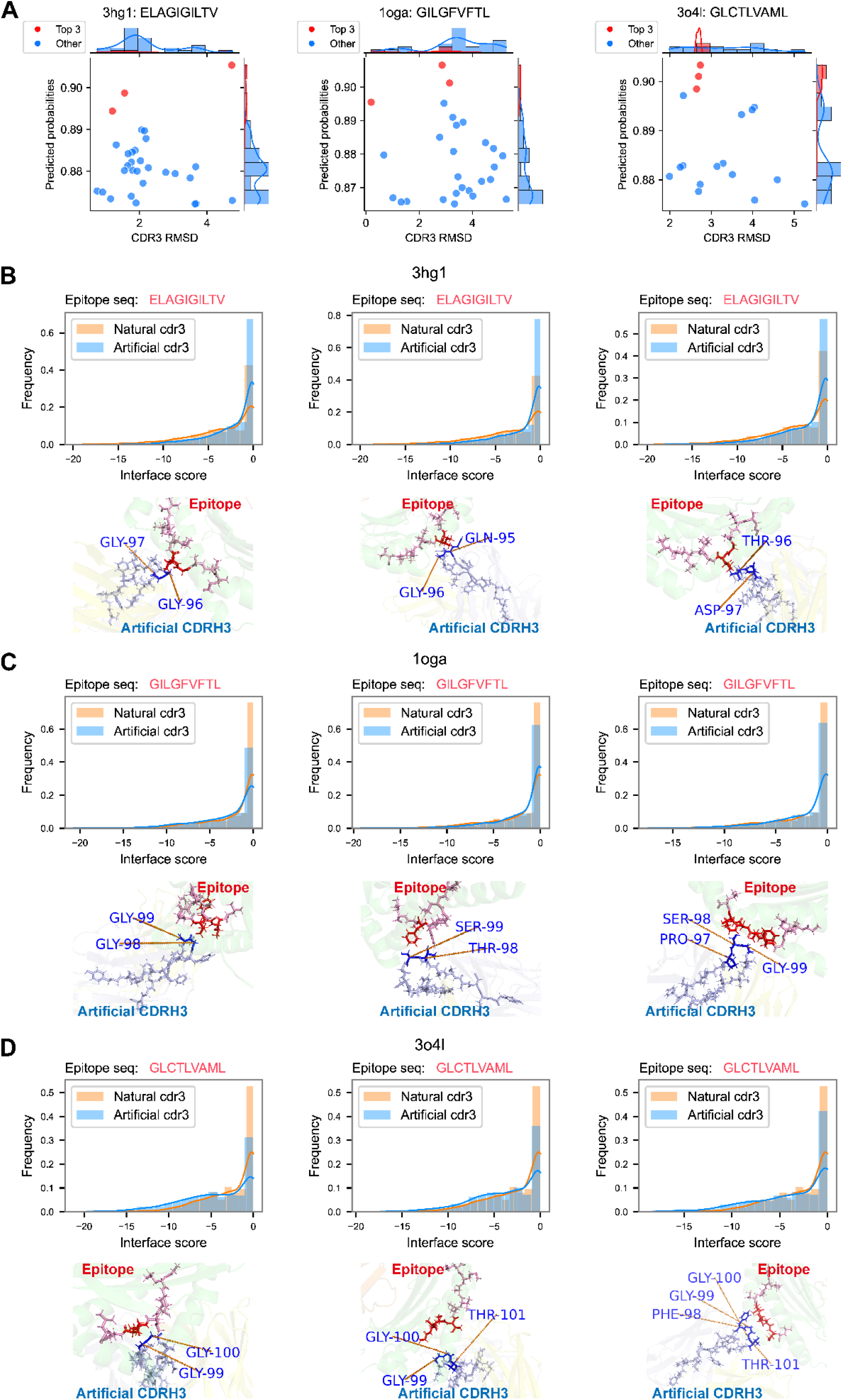
The comparison of the generated TCRs with the natural ones in view of their structural similarity and binding affinity. We selected three epitopes “ELAGIGILTV” (PDB ID: 3hg1), “GILGFVFTL” (PDB ID: 1oga), “GLCTLVAML” (PDB ID: 3o4l) and their corresponding natural TRBs “CAWSETGLGTGELFF” (3hg1), “CASSSRSSYEOYF” (1oga), “CSARDGTGNGYTF” (3o4l) for which real structures are available at PDB (protein data bank). A. The relationship between the estimated root-mean-square deviation (RMSD) with the predicted probabilities generated by ERTransformer. Each dot denotes one artificial TRB, with those having the top 3 predicted probabilities highlighted in red. B-D. The estimated interface scores of TRBs towards corresponding epitopes (B-3hg1, C-1oga, D-3o4l). The bar plots in B-D present the distribution of the interface scores, with natural and artificial TRBs highlighted in orange and blue, respectively. The estimated optimal docking positions are presented in the lower figures in B-D, with the MHC and TCR regions represented in green and yellow, respectively. The nearest amino acid positions (such as GLY-96 and GLY-97 in B) are highlighted in B-D.

Therefore, we further investigated their binding affinity for the top 3 artificial TRB sequences (Figure 4A). This analysis was based on the interface score of the binding complex between TRBs and the pMHC, using RosettaDock-4.0 ^27^. As presented in the histogram in Figure 4(B, C, D), the top 3 artificial TRBs exhibit similar or even lower interface scores compared with the natural TRBs (CDR3 part), visualized CDR3 and epitope structure also showed close positions. These results indicate that the generated artificial TRBs present a binding affinity towards their corresponding epitopes that closely resembles that of natural TRBs.

### 2.6 ERTransformer identifies key amino acid positions crucial for TCR-pMHC docking determination

The attention layers are a key component of EpitopeBERT and ReceptorBERT, allowing these two BERT models to focus on certain amino acid tokens in a sequence and weigh them more heavily in their analysis, rather than considering all tokens equally. To validate whether the ERTransformer model constructed by ER-BERT captures the binding rules between epitopes and TCRs, we investigated a number of key amino acid positions using the integrated gradients method ^28^ and compared these positions with the actual contact residues. We collected crystallography data from The Protein Data Bank (PDB) for epitopes “GILGFVFTL” (PDB ID: 1oga) and “GLCTLVAML” (PDB ID: 3o4l), two epitopes that also existed in the 10X dataset. To visualize the attribution scores of each amino acid in the epitope to the generated TRB, we used ER-BERT with the UAA tokenizer (each token represents an amino acid) after fine-tuning the Seq2Seq task (ERTransformer). For a given amino acid in the epitope, its attribution scores indicate the importance of its contribution in determining the amino acids in the receptor. A positive value indicates a positive contribution, while a negative value indicates a negative contribution.

Figure 5(A) and (B) present the heatmap of attribution scores for the epitopes “GILGFVFTL” and “GLCTLVAML” respectively, with red indicating a higher positive weight and blue indicating a higher negative weight. We also created a sequence motif plot according to the average attribution scores of all the amino acids in the generated TRBs for each amino acid in the two epitopes, as shown in Figure 5(C) and (D). Our analysis revealed that the ERTransformer tended to focus on the central regions of these two epitopes. Particularly, for “GILGFVFTL”, ERTransformer focused on the third and seventh amino acids, that is Leucine (L) and Phenylalanine (F). For “GLCTLVAML”, ERTransformer assigned higher weights to the second, third, fifth, and sixth amino acids, which are Leucine (L), Cysteine (C), Leucine (L), and Valine (V). However, when examining the TRB sequences, we did not observe clear trends. This might be because, during the training in the Seq2Seq task, the ERTransformer model was fed with epitopes to generate TRB sequences, but did not directly use TRB sequences as input. The crystallography structures of two epitopes with the corresponding CDR3 part of TCR (both TRA and TRB), and HLA or MHC and their relative positions are shown in Figure 5(E) and (F). We found that the amino acid positions with higher weights are highly likely to be congruent with the amino acids determining the folding positions, which subsequently or directly determined known important docking positions ^40–42^. For instance, in the epitope “GILGFFTL”, the first Leucine (L) determines the folding positions of this epitope, and the second phenylalanine (F) is the closest position to the TRB (green) binding site ^41^. Similarly, in the epitope “GLCTLVAML” the first Leucine (L) and Cysteine (C) determine the first folding positions, and the second Leucine (L) and Valine (V) are also the closest positions to the TRB (green) binding site ^40^. To summarize, through the attention layers, ERTransformer could identify some key amino acid at specific positions, which may determine the binding between epitopes and TCRs.

**Figure 5.**
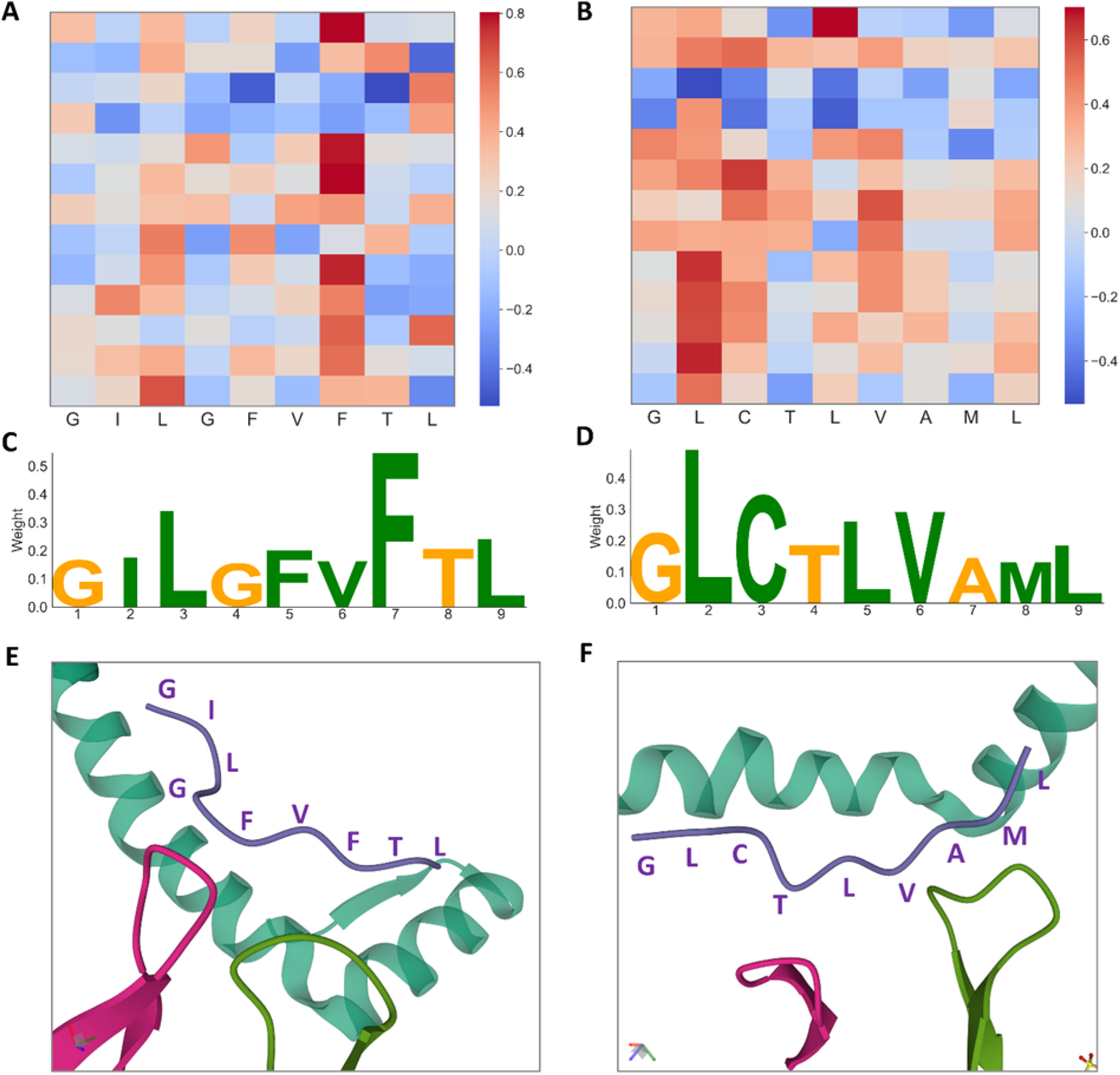
ERTransformer can capture the amino acid positions that determine the docking positions. We used two epitopes “GILGFVFTL” (PDB ID: 1oga) and “GLCTLVAML” (PDB ID: 3o4l) as examples here. These two epitopes have the crystallography data from The Protein Data Bank and also existed in our external dataset - 10X. (A) “GILGFVFTL” and (B) “GLCTLVAML” show the heatmap of attribution weights for each amino acid computed by integrated gradients on the ERTransformer with the UAA tokenizer (each token represents an amino acid), with red indicating a higher positive weight and blue indicating a higher negative weight. (C) and (D) are the sequence motif plots of “GILGFVFTL” and “GLCTLVAML”, respectively. The weights in (C) and (D) come from the average attribution scores of all the amino acids in the generated TRBs for each amino acid in the two epitopes. (E) and (F) show the visualization of the crystallography structures of two epitopes (purple) with the corresponding CDR3 parts of TRA (red) and TRB (green), and HLA or MHC (dark green) and their relative positions. Note that we only keep the structures that are closest to the epitope for better visualization.

## 3. Discussion

In this paper, we present ERTransformer for generating TCR (TRB) sequences with the desired epitope-binding properties. ERTransformer was constructed from ER-BERT, which consists of EpitopeBERT and ReceptorBERT. We first demonstrated ER-BERT could learn biological meaningful representations of epitopes and receptors by introducing them to cluster epitope-specific TCRs. Then, we validated that ERTransformer could generate artificial TCRs that have low sequence and structure identity, while concurrently exhibiting a marked similarity in biological functionality and binding affinity to their natural TCR counterparts. The artificial TCRs demonstrated great binding specificity to the corresponding epitopes. Additionally, we found ERTransformer could capture the key amino acids that determine the docking positions of epitopes and TCRs. ERTransformer is a data-driven tool for studying the immune system and for developing advanced immunotherapies. By generating high-quality TCR sequences and identifying key amino acids, ERTransformer can help researchers better understand the mechanisms of immune recognition and response.

The CDR3 region, despite being a fraction of the full-length sequence, contributes substantially to the diversity of the TCR repertoire and is vital for antigen specificity. While the complete TCR structure, including CDR1 and CDR2 region, as well as the TCR alpha chain, also plays a crucial role in TCR binding, ER-BERT’s capability to generate TCR sequences, particularly those of the CDR3 region, holds considerable value. It serves as a robust resource for initial computational investigations and contributes to efforts in predictive modeling. We demonstrated that the learned representations of ER-BERT are biologically meaningful, which can provide superior epitope-specific clustering of TCRs. The ER-BERT-BSP is specially designed for binding specificity prediction while using the learned representations from the epitope and TCR simultaneously. The superior performance of ER-BERT-BSP compared to DeepTCR, which only uses representations of TCRs, indicates that the addition of the epitope sequence to the model improves the learning of binding rules between the epitope and TCR. Epitopes are an essential part of the interaction and considering their sequences allows ER-BERT-BSP to capture a fuller picture of the binding rules. Furthermore, ER-BERT-BSP employs the sophisticated BERT architecture, designed specifically to master the contextual representation of sequences. This contrasts with DeepTCR, which utilizes convolutional neural networks and variational auto-encoder structures, neither of which are explicitly designed to capture the intricate dependencies found in sequence data. This architectural divergence likely contributes to the performance differential observed between ER-BERT-BSP and DeepTCR.

Recent advances in single-cell multi-omics have provided an opportunity to train deep-learning models using large amounts of sequence data to address the challenging task of TCR generation. Our results show that ERTransformer significantly outperforms Semi-Random-Transformer, which is composed of the latest model on the TCR-related task (TCRBert). The superior performance of ERTransformer comes from two modeling mechanisms: First, ERTransformer utilizes the most comprehensive epitope and TRB sequences so far, including tens of thousands of epitope and TRB sequences from 12 datasets; Second, ERTransformer utilizes the framework of pre-training and fine-tuning and specifically designed for sequence generation task. The epitope blocks and receptor blocks used in ERTransformer for the TCR sequence generation task are inherited from the pre-trained ER-BERT model on the MAAT task, which contains rich knowledge of the rules existing in epitope and TCR sequences. Compared with the baseline that combines a randomly initialized BERT with TCRBert (Semi-Random-Transformer), this rich knowledge helps ERTransformer better capture the binding rules from epitopes to receptors to better generate epitope-binding TCR sequences. In addition, considering the performance on the binding specificity prediction task, ER-BERT-BSP still significantly outperforms the latest deep learning models such as DeepTCR. As a model uses the sequence data only, we do not have to feed any other biological information, such as VDJ gene usage in DeepTCR, to ER-BERT-BSP, but with the advantage of pre-training and a large amount of sequence data, ER-BERT-BSP still outperforms other state-of-the-art models.

Despite the promising results of our study, ERTransformer has some limitations that need to be addressed in future work. ERTransformer currently uses epitope sequences to generate receptor sequences, but can not control the binding affinity between epitopes and the generated receptors. In future work, we plan to investigate improved strategies to generate TCRs with desired binding affinities. Additionally, due to the limited data of TRA sequences, ERTransformer currently is only applicable to TRB sequences. In the future, we plan to extend ERTransformer to other types of immune system sequences, such as TCR alpha or B cell receptor sequences. Moreover, the validation of ERTransformer is only *in silico* experiments. We plan to experimentally assess the quality of the generated TCR sequences by conducting *in vitro* experiments in future work.

In this study, taking the generation of epitope-binding TCR as an example, we demonstrate the feasibility of generating interacting partner proteins according to the sequence of a given protein. It’s now possible to construct diverse generative models for the generation of artificial proteins with desired binding properties to a target protein. These artificial proteins would be powerful biomedical and bioengineer tools to regulate biological processes, treat diseases and even form new biological functions.

## 4. Methods and Datasets

ERTransformer consists of the epitope (encoder) and receptor (decoder) blocks derived from ER-BERT, i.e. EpitopeBERT and ReceptorBERT, which both utilize a commonly-used BERT architecture. BERT^18^ was initially proposed to deal with the natural language processing tasks, such as translation ^29^, and sentiment analysis^30^. The BERT model consists of 12 encoder blocks, and each encoder block consists of two layers: multi-head self-attention layers and feed-forward layers. Each multi-head self-attention layer has 12 heads. Given an input sequence, instead of using the embeddings of each token (word) directly, the self-attention module would create three vectors called *query (Q)*, *key (K)*, and *value (V)* and then compute attention scores to get a better encoding for each token. The output of the self-attention module is then fed into the feed-forward layer to the next encoder block. The architecture of stack encoder blocks could allow the model to learn high-dimensional and complex interactions between tokens that better capture the “grammar” rules that existed in the input sequences. For more details on BERT models, please refer to the original paper ^18^.

### 4.1 Tokenizer

Unlike English sentences, which have natural spaces between words, epitope and TCR sequences are amino acid sequences without any separation. And the basic processing unit in the BERT model is a token, or word. To address this issue, we developed two different methods inspired by Forward Maximum Matching ^43^, which is used in Chinese language processing. These two tokenizers are described in detail below:

#### 4.1.1 Unique Amino Acid (UAA) Tokenizer

As a common practice, the UAA tokenizer treats each amino acid as a token.

#### 4.1.2 Forward Maximum Frequency Matching (FMFM) Tokenizer

Motifs, a short conserved amino acid sequence pattern, are widely present in many proteins ^31^, and their frequencies are highly associated with distinct functions ^32^. FMFM tokenizer is based on an assumption that the motifs with high frequencies will be more important for their corresponding protein’s biological functions ^44,45^. As shown in Figure 1(B), the details of the FMFM tokenizer are as follows: Get the motifs with specific lengths. For all the sequences *S* belonging to the same category (epitope, TRA, and TRB), we split each sequence into motifs with a specific length *L* (such as 3). That is, the next motif will start with the position following the previous one. For example, given the epitope sequence “ATDALMTGY”, the split motifs with length 3 are “ATD”, “ALM”, “TGY”. In this study, we consider motifs with length 2 and length 3.

1. Build the motif vocabulary. After obtaining all motifs of a certain length from the sequences belonging to the same category, we count their frequencies and use z-score normalization to make fair comparisons between motifs of different lengths.
2. Tokenization. Given one sequence, the FMFM tokenizer will consider both the length and frequency of specific motifs. For example, given the epitope sequence “ATDALMTGY”, the FMFM tokenizer will first consider the frequencies of motif “AT” and “ATD”, and suppose their normalized frequencies are 0.3 and 1.2, respectively, FMFM tokenizer will get the second token whose normalized frequency is the largest, that is “TD”. The next

token will start with the position following the previous one - “ATD”, that is starting from “ALMTGY”, and repeat this process to the end of the sequence. One possible scenario is that the end of the sequence after this process will be a single amino acid, which we will keep as the last token directly (Figure 1(B)).

### 4.2 Training of ER-BERT

The general framework of ER-BERT is shown in Figure 1(C-E). Given the input epitope or TCR sequence with length *l*, after tokenization, the input is *M* tokens. These tokens are then padded with two special tokens as a common approach used in BERT models: a classification token *[CLS]* as a prefix and a separator token *[SEP]* as a suffix. Then, the *M* + 2 tokens will be fed into an embedding layer to encode each token to a continuous representation of 768 dimensions. Adding the positional embeddings together, the summed sequence embedding is fed into EpitopeBERT or ReceptorBERT which consists of 12 encoder blocks. After that, we input the sequence embedding after the BERT model to a “head” layer for certain pre-training or downstream tasks.

#### 4.2.1 Masked Amino Acid Token (MAAT) Task

Following the Masked Language Model (MLM) task commonly used in BERT models, we pre-train both EpitopeBERT and ReceptorBERT using a Masked Amino Acid Token (MAAT) Task. As shown in Figure 1(C), we randomly hide or mask 15% of the tokens in each input epitope or TCR sequence, and train EpitopeBERT and ReceptorBERT to predict the masked tokens (denoted as *[MAAT]*). We append an “MAAT” head to both EpitopeBERT and ReceptorBERT as the last output module, which is a two-layer fully connected neural network connected with the Gaussian Error Linear Unit (GELU) activation function ^46^. The output dimension of the “MAAT” head is consistent with the number of unique tokens, followed by a softmax activation function. MAAT task is performed on all the available epitope and TCR sequences to learn the basic “grammar” rules existing in these sequences.

#### 4.2.2 Binding Specificity Prediction (BSP) Task

To allow ER-BERT to capture the binding specificity rules of epitopes and TCRs, we connect EpitopeBERT and ReceptorBERT and train these two BERT models together on the binding specificity prediction task. As shown in Figure 1(D), given one epitope-TCR pair, we feed the epitope and TCR to the EpitopeBERT and ReceptorBERT separately and get the embedding of the classification token *[CLS]* for the following operation. We append a “BSP” head to get the final output, which is a three-layer fully connected neural network connected with tanh activation function. The final output is one value through softmax activation, denoting the probability of whether the input epitope could bind to the TCR. The BSP task aims to combine EpitopeBERT and ReceptorBERT together and allow ER-BERT to master the “binding rule” of epitopes and TCRs.

In addition, since most datasets only provide the positive epitope-TCR pairs, which indicate that a TCR binds to an epitope, there are no negative pairs in our dataset. To deal with the scarcity of negative epitope-TCR pairs, we developed a negative sampling method for the BSP task. The fundamental assumption here is that given one epitope, the probability of one randomly-selected TCR being incompatible to bind to the epitope is much greater than the probability of affinity. The negative sampling method consists of two steps:

1. For each epitope, suppose we have *n*_p_ positive TCRs that can bind to it, we sample *n*_p_ TCRs from all the TCR sequences as its corresponding negative TCRs. Note that these negative TCRs are chosen not only from the TCRs that do not form positive pairs with the specific epitope, but also from TCRs that form positive pairs with other epitopes.
2. To enrich the diversity of epitopes in the BSP task, we randomly select *n* epitope-TCR pairs as negative samples from all possible pairs, ensuring that these pairs have not been identified as positive. This strategy broadens the range of epitopes and TCRs in the negative sample set, while maintaining the condition that no epitope-TCR pair in the negative set has been identified as positive.

#### 4.2.3 Seq2Seq Task

Following the idea that utilizes two BERTs as Encoder and Decoder separately to build a Transformer model ^47,48^, we use EpitopeBERT to build epitope blocks as Encoder and ReceptorBERT to build receptor blocks as Decoder to form a Transformer called ERTransformer for the Seq2Seq training (Figure 1(E)). That is, given the epitopes, we train the ERTransformer to generate corresponding TCRs. Specifically, except that the parameters of the cross-attention layer in the receptor block are initialized randomly, the rest parameters of the epitope block and receptor block are inherited from EpitopeBERT and ReceptorBERT, respectively. During the training of the Seq2Seq task, the encoder layer will be fed the whole sequence of the epitope. The decoder will generate the target TCR sequences step by step. At each step, the input to the decoder layer is the previously predicted tokens of TCR concatenated with the to-be predicted token (masked as [pad]). As shown in the expression below, the *key* (*Q_E_*) and *value* (*K_E_*) matrix generated by the epitope block are communicated with the *query* (*V*_*R*_) matrix from the receptor block in the cross-attention layer, where *Z* is the output of the cross-attention layer and *d_k_* is the dimension of the embeddings.

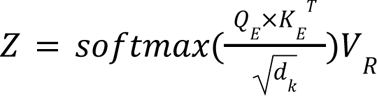

Then, we utilized the cross-entropy loss computed between the predicted token with the true one.

### 4.3 Fine-tuning of ERTransformer on a new dataset

To utilize ERTransformer for the TCR generation for specific epitopes, the primary issue is how to determine the quality of the generated TCRs. In this study, we train an external discriminator based on the BSP task. In detail, as shown in Figure 1(D), given a dataset that contains the natural epitope-TCR pairs, for each epitope, we use 80% of its binding TCR sequences (named *D*_*d*_) to train the external discriminators and the left 20% TCR sequences (named *D*_*g*_) are used to fine-tune the trained ERTransformer on the Seq2Seq task. This separation of the dataset aims to ensure that the generator and discriminator do not overlap, and maximize the performance of both at the same time. The utilization of ERTransformer for TCR generation is as follows:

1. Fine-tune ERTransformer. To allow the ERTransformer better capture the patterns of amino acids of the binding TCRs of one epitope, we first fine-tune the trained ERTransformer on *D*_*g*_. The TCRs in *D*_*g*_ perform as “seed” TCRs for the following TCR generation.
2. Train external discriminator. The external discriminator can be the trained ER-BERT after the BSP task, or some state-of-the-art models on this task, such as DeepTCR ^35^. DeepTCR was originally developed as a multi-class classifier given a set of TCRs. In this study, we found that DeepTCR’s original version achieves a lower performance on *D*_*g*_ regardless of MIRA or 10X, thus we utilize DeepTCR to train one binary classifier for each epitope separately for better performance.
3. Determine the quality of the generated TCRs. We use the beam search ^33^ method to generate new TCRs. Beam search is a heuristic algorithm and is widely used in many translation tasks. We use the fine-tuned ERTransformer to generate 1,000 TCRs for each epitope and utilize external discriminators to determine how many generated TCRs can bind to the input epitope.

### 4.4 Datasets for training

We collected the sequences of epitopes and CDR3 sequences of TCRs (TRA and TRB) from 9 public datasets and 2 in-house datasets. Among all the datasets, we only kept valid epitopes and TCRs that contain 20 standard amino acids from human species. All these sequences and epitope-TCR pairs were used for the training of MAAT, BSP, and Seq2Seq tasks. The details of each dataset are shown in Table 1. We found that the number of TRB sequences is far larger than that of TRA sequences, thus we focus on TRB sequences in this study. In general, we collected 1,929,016 epitope sequences and 33,088,640 TRB sequences for the MAAT task for EpitopeBERT and ReceptorBERT, respectively. For the BSP and Seq2Seq tasks, we prepared 176,268 positive epitope-TRB pairs.

**Table 1.**
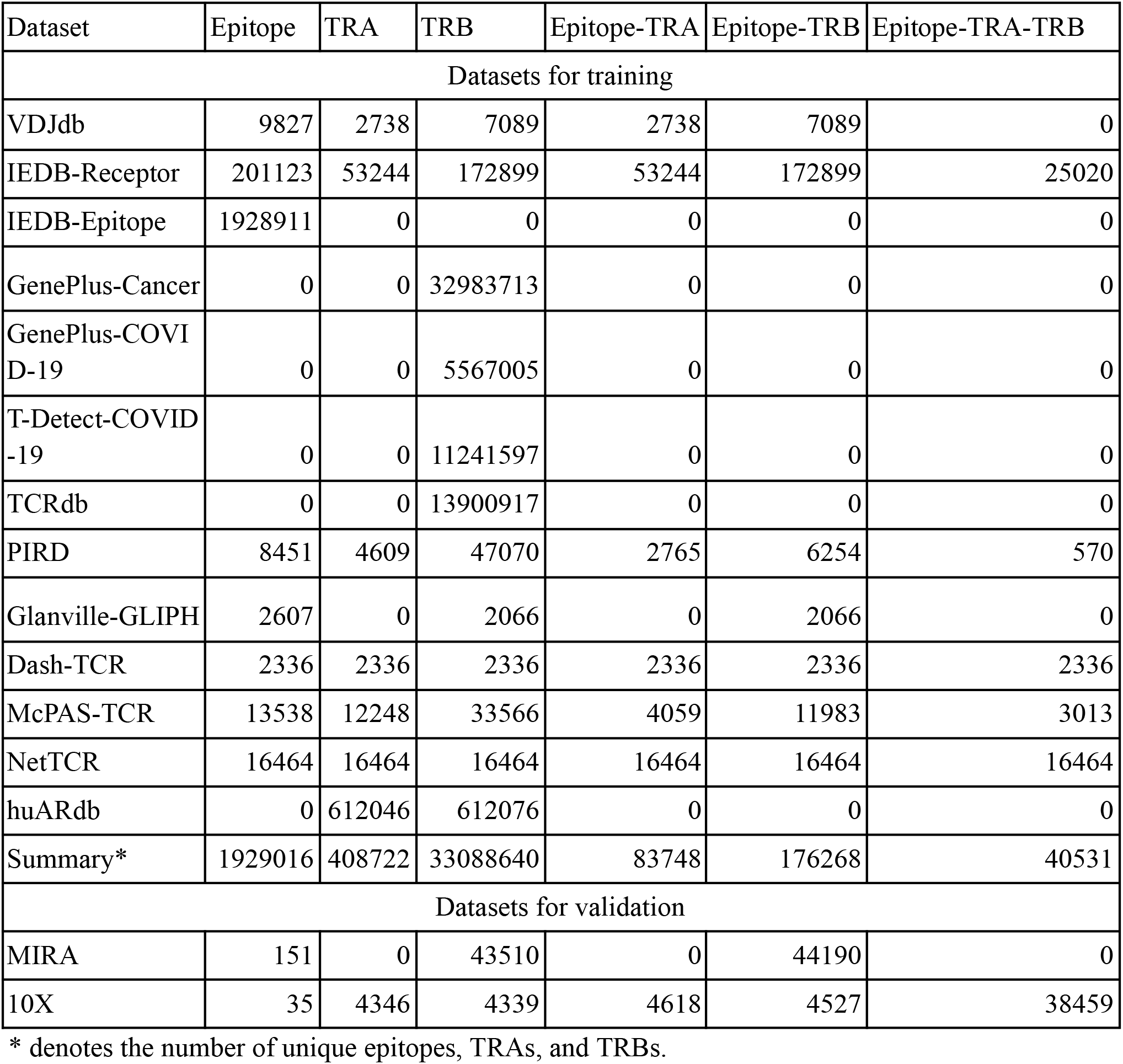
The statistics of valid epitopes, TCRs, and their pairs.

| Dataset | Epitope | TRA | TRB | Epitope-TRA | Epitope-TRB | Epitope-TRA-TRB |
| --- | --- | --- | --- | --- | --- | --- |
| Datasets for training |  |  |  |  |  |  |
| VDJdb | 9827 | 2738 | 7089 | 2738 | 7089 | 0 |
| IEDB-Receptor | 201123 | 53244 | 172899 | 53244 | 172899 | 25020 |
| IEDB-Epitope | 1928911 | 0 | 0 | 0 | 0 | 0 |
| GenePlus-Cancer | 0 | 0 | 32983713 | 0 | 0 | 0 |
| GenePlus-COVID-19 | 0 | 0 | 5567005 | 0 | 0 | 0 |
| T-Detect-COVID-19 | 0 | 0 | 11241597 | 0 | 0 | 0 |
| TCRdb | 0 | 0 | 13900917 | 0 | 0 | 0 |
| PIRD | 8451 | 4609 | 47070 | 2765 | 6254 | 570 |
| Glanville-GLIPH | 2607 | 0 | 2066 | 0 | 2066 | 0 |
| Dash-TCR | 2336 | 2336 | 2336 | 2336 | 2336 | 2336 |
| McPAS-TCR | 13538 | 12248 | 33566 | 4059 | 11983 | 3013 |
| NetTCR | 16464 | 16464 | 16464 | 16464 | 16464 | 16464 |
| huARdb | 0 | 612046 | 612076 | 0 | 0 | 0 |
| Summary* | 1929016 | 408722 | 33088640 | 83748 | 176268 | 40531 |
| Datasets for validation |  |  |  |  |  |  |
| MIRA | 151 | 0 | 43510 | 0 | 44190 | 0 |
| 10X | 35 | 4346 | 4339 | 4618 | 4527 | 38459 |

\* denotes the number of unique epitopes, TRAs, and TRBs.

#### VDJdb

VDJdb is a curated database of TCR sequences with known epitope specificities ^49^. The TCRs in VDJdb come from human, mouse, and macaque. We kept the data that belong to humans and with confidence scores larger than 0.

#### IEDB

IEDB^50^ consists of the Receptor (IEDB-Receptor) dataset and Epitope (IEDB-Epitope) dataset and contains both TCRs and BCRs. We only keep TCR sequences for the IEDB-Receptor dataset. The IEDB-Epitope dataset only contains epitopes and 1,928,911 epitopes left after curation.

#### T-Detect-COVID

This dataset is generated by the T-Detect COVID Test which can detect the specific T cell signature (TRB) in response to SARS-CoV-2 and only contains TRBs. The t-Detect-COVID dataset is made public by Adaptive Biotechnologies and Microsoft ^51^.

#### TCRdb

TCRdb^52^ is a comprehensive TCR database that mainly focuses on TRB sequences without epitope specificities.

#### PIRD

Pan immune repertoire database (PIRD)^53^ collects raw and processed TCRs of human and other vertebrate species.

#### Glanville-GLIPH

Glanville et al.^15^ developed an algorithm GLIPH to cluster TCRs (TRBs). We collected 2,607 epitopes and 2,066 TRBs from the data they used to develop GLIPH.

#### Dash-TCR

Dash et al.^54^ used molecular genetic tools to analyze the diversity of epitope-specific TCRs. We collected 2446 epitopes, TRAs and TRBs from the data this study used.

#### McPAS-TCR

McPAS-TCR^55^ is a manually curated catalogue of TCRs that existed in humans and in mice. We collected 13,538 epitopes, 12,248 TRAs, and 33,566 TRBs from the curated dataset.

#### NetTCR

NetTCR is an algorithm developed by Morten et al.^56^ which enables accurate prediction of TCR-peptide binding. We collected 16,464 epitopes, 16,464 TRAs, and 16464 TRBs used in NetTCR.

#### GenePlus

We have two in-house TRB bulk-sequencing datasets collected for studying Cancer and COVID-19, respectively. The cancer data includes 32,983,713 TCRBs from over 1000 cancer patients. The COVID-19 data includes 5,567,005 TCRBs from 48 participants at different time points before and after receiving the COVID-19 vaccine. Geneplus Co Ltd produced these two datasets.

#### huARdb

The huARdb database is a large-scale human single-cell immune profiling database that contains 612,046 high-confidence T or B cells with full-length TCR/BCR sequences and transcriptomes from 215 datasets ^57^.

### 4.5 Datasets for validation

In this study, we test and validate the performance of ER-BERT on two external TCR datasets. **MIRA.** MIRA dataset is about the TRB sequences from subjects exposed to or infected with the SARS-CoV-2 virus. The original MIRA dataset includes more than 135,000 high-confidence SARS-CoV-2-specific TCRs. And after curation, we have 154 epitopes and 44,190 epitope-TRB pairs.

**10X.** We downloaded the epitope-specific binding data of paired TCRs from the 10x Genomics website (https://support.10xgenomics.com/single-cell-vdj/datasets). After data curation, 38,459 paired TCR TRA and TRB chains that bind to 35 epitopes, including GILGFVFTL from the M1 protein of the influenza virus (flu), IVTDFSVIK from the Epstein-Barr nuclear antigen 4 of the Epstein-Barr virus (EBV), LTDEMIAQY from the SARS-CoV-2 Surface GP 1 protein of the SARS-CoV-2 virus, and so on.

### 4.5 Evaluation methods

#### 4.5.1 Clustering and DBI scores

The initial embedding of each epitope or TCR generated by ER-BERT is represented in 768 dimensions. In order to generate visual plots, we conducted Principal Component Analysis (PCA) using the *sklearn.decomposition.IncrementalPCA* module, resulting in a 50-dimensional representation of epitopes and TCRs. Subsequently, we utilized t-SNE (implemented in *sklearn.manifold.TSNE*) to obtain 2-dimensional representations for visualization and DBI calculation. With the 2-dimensional representations of all epitopes and TCRs generated by the model, we employed *sklearn.metrics.davies_bouldin_score* to compute the DBI scores. Note that the input *X* in the function *sklearn.metrics.davies_bouldin_score* corresponds to the 2-dimensional representations, while *Y* represents the target epitopes for each generated TCR.

#### 4.5.2 The retraining of external discriminators - DeepTCR

DeepTCR, initially developed by Sidhom et al. ^35^, was utilized in this study for the classification of antigen-specific TCRs. To adapt DeepTCR for the 10X and MIRA datasets, we employed the same training methodology as ER-BERT-BSP. Specifically, DeepTCR was employed to construct a classifier for each epitope. In line with this approach, we utilized 80% of the data to retrain a single DeepTCR classifier that encompassed all epitopes within the 10X and MIRA datasets. The source code for DeepTCR, which was employed in this study, can be accessed at https://github.com/sidhomj/DeepTCR.

#### 4.5.3 The fine-tuning of Semi-Random-Transformer

The Semi-Random-Transformer is created based on the latest BERT model TCRBert ^19^ for the modeling of the grammar existing in TCR sequences. Specifically, we created a Transformer model using function *transformers.EncoderDecoderModel.from_encoder_decoder_pretrained(encoder_pretrained_mo del_name_or_path, decoder_pretrained_model_name_or_path)*, where *encoder_pretrained_model_name_or_path* is the path of pre-trained TCRBert (available at https://huggingface.co/wukevin/tcr-bert), and *decoder_pretrained_model_name_or_path* denotes the path of a randomly initialized BERT without any prior training). The fine-tuning process for the Semi-Random-Transformer aligns with that of the ERTransformer, as elaborated in Section 4.3 of this paper.

#### 4.5.4 The sequence similarity measure methods: BLAST and BLOSUM62 matrix

The Basic Local Alignment Search Tool (BLAST) ^22^ tool is used to compare its amino acid sequence and calculate the statistical significance of the matched reference sequence. In this study, we built a reference sequence dataset using all the TCR sequences presented in the MIRA and 10X except the sequences used to fine-tune the models on the Seq2Seq task. Given all the TCR sequences generated by one model for one epitope, we utilized the command *blastp -query epitope.fasta -db db10x -out epitope.txt -task blastp-short -outfmt “6 qseqid sseqid sseq evalue bitscore pident positive” -num_threads 10* to get the bit-scores. For each generated TCR sequence that was also determined to bind to the given epitopes, we kept the matched reference TCR sequence with an e-value smaller than 0.001 and recorded the bit-score. Bit-score is used to determine the sequence similarity in the BLAST tool, and the higher the bit-score, the better the sequence similarity.

The BLOSUM62 matrix used in this study is available at https://resources.qiagenbioinformatics.com/manuals/clcmainworkbench/current/index.php?manu <u>al=BE_Scoring_matrices.html</u>. For one model, we retrieved all one epitope’s generated TCRs and natural ones. To compute BLOSUM62 score, we aligned all these TCR sequences by keeping sequences with the maximum longest length. For one amino acid *i* in the generated TCR sequence and its corresponding amino acid *j* at the same position in another sequence, we obtained the BLOSUM62 score for that position by referencing the value in the BLOSUM62 matrix, denoted as *BLOSUM62_ij_*. Then, we iteratively calculated the sum of scores by traversing through each position.

#### 4.5.5 The structure similarity and binding affinity by existing computational docking methods

Given one generated artificial TCR and one natural TCR, their structure similarity of the CDR region is measured by the root-mean-square deviation (RMSD) of atomic positions. Specifically, due to the unavailability of the generated artificial TCR’ structure, we obtained the complete sequence of the corresponding natural TCR and substituted the original CDR3 segment with the generated CDR3 region. Subsequently, we introduced TCRdock ^26^ to predict the structures of artificial TCRs. Then we used PyMOL to visualize the structure. The RMSD for the CDR3 region is computed by PyMOL by specifically selecting the CDR3 region.

To investigate the binding affinity between the generated artificial TCRs and epitopes, given the estimated structure of generated artificial TCRs and experimental validated structure of epitopes available at PDB, we first employed TCRDock ^26^ to estimate the correct docking positions between (top-3 in Figure 4(A)) and the epitopes in the pMHC. Then, we introduced RosettaDock-4.0 ^27^, using the previously determined docking positions as initial coordinates, to predict the binding affinity. The binding affinity is measured by interface scores, where a smaller indicates a stronger binding affinity. Specifically, we utilized the server version of RosettaDock-4.0, which is available at https://r2.graylab.jhu.edu/apps/submit/docking. RosettaDock-4.0 was employed to perform 1,000 experiments with different potential docking positions, resulting a distribution of interface scores as presented in Figure 4.

## 5. Data availability

All described public datasets are available through the corresponding repositories. VDJdb: https://vdjdb.cdr3.net; IEDB: https://www.iedb.org/; T-Detect-COVID: https://www.adaptivebiotech.com/immunecode/; TCRdb: http://bioinfo.life.hust.edu.cn/TCRdb/<u>#/</u>; PIRD: https://db.cngb.org/pird/home/; Glanville-GLIPH: https://github.com/immunoengineer/gliph; Dash-TCR: https://www.nature.com/articles/nature22383#Sec22; McPAS-TCR: http://friedmanlab.weizmann.ac.il/McPAS-TCR/; NetTCR: https://github.com/mnielLab/NetTCR-2.0; huARdb: https://huarc.net/database; MIRA: https://clients.adaptivebiotech.com/pub/covid-2020; 10X: https://www.10xgenomics.com/resources/datasets.

## 6. Code availability

The source codes of ER-BERT are available at https://github.com/TencentAILabHealthcare/ER-BERT. The pre-trained EpitopeBERT and ReceptorBERT model files, and the fine-tuned ER-BERT-BSP and ERTransformer model files are available at https://doi.org/10.5281/zenodo.7494046.

